# Potency of CRISPR-Cas Antifungals Is Enhanced by Co-targeting DNA Repair and Growth Regulatory Machinery at the Genetic Level

**DOI:** 10.1101/2023.07.11.548496

**Authors:** Brian J. Mendoza, Xianliang Zheng, Jared C. Clements, Christopher Cotter, Cong T. Trinh

## Abstract

Fungal pathogens are virulent and resistant to antibiotic treatment. Due to their adaptability, specificity, and efficiency, CRISPR-Cas systems can be harnessed for neutralizing these fungal pathogens. However, the conventional design of CRISPR-Cas antimicrobials, based on induction of DNA double-strand-breaks (DSBs), is potentially less effective in fungi due to robust eukaryotic DNA repair machinery. Here, we report a novel design principle to formulate more effective CRISPR-Cas antifungals by co-targeting essential genes with DNA repair defensive genes that remove the fungi’s ability to repair the DSB sites of essential genes. By evaluating this design on the model fungus *Saccharomyces cerevisiae*, we demonstrated that essential and defensive gene co-targeting is more effective than either essential or defensive gene targeting alone. The top-performing CRISPR-Cas antifungals performed as effectively as the antibiotic Geneticin. Fast growth kinetics of *S. cerevisiae* induced resistance to CRISPR-Cas antifungals where genetic mutations mostly occurred in defensive genes and guide RNA sequences.

**Significance:** The emergence of virulent, resistant, and rapidly evolving fungal pathogens poses a significant threat to public health, agriculture, and the environment. Targeting cellular processes with standard small-molecule intervention may be effective but requires long development times and is prone to antibiotic resistance. To overcome the current limitation of antibiotic development and treatment, this study harnesses CRISPR-Cas systems as antifungals by capitalizing on their adaptability, specificity, and efficiency in target design. Simultaneous co-targeting of both essential and defensive genes is shown to be a novel design principle for formulating effective CRISPR-Cas antimicrobials that can be rapidly tuned to adapt to inevitable escapee events.

## Introduction

Fungi represent a particularly resistant class of pathogens that are the source of dysbiosis in a variety of societally relevant hosts and environments (1, 2). In humans, fungal pathogens are responsible for over one billion infections and 1.5 million deaths each year with species of *Candida, Aspergillus,* and *Cryptococcus* accounting for 90% of fungal-related deaths (3). The prevalence of fungal infections is surging due to an increase in immunocompromised individuals, as well as an increase in geographic range and dispersal due to climate change (4, 5). Along with an increased at-risk population, the threat of fungal pathogens is amplified by the rapid emergence of antifungal resistance. Overuse of current antifungals in both medical and agricultural settings has escalated the emergence of antifungal-resistant species (6). Azole resistance is widespread in *Candida, Aspergillus,* and *Cryptococcus spp.*, while echinocandin and polyene resistance is less frequent, but has been found in several *Aspergillus*, *Candida,* and *Cryptococcus* species (6, 10, 11). Most notably, *Candida auris,* a fungal pathogen first identified in Japan in 2009 that since has spread globally with cases rising in parallel with the COVID-19 pandemic, is often multidrug resistant with 90% of isolates being fluconazole resistant and (7–9). The increasing threat of antifungal-resistant species and the lack of new antifungal drugs warrants the investigation into novel antifungal strategies.

CRISPR-Cas (clustered regularly interspaced short palindromic repeats-CRISPR– associated) gene editing, a powerful method for knocking out key genes in the growth regulation cycle, has recently shown promise for antimicrobial treatment in bacterial systems (12–14). Adaptability, specificity, and efficiency in designing guide RNAs (gRNAs) to enable precise gene inactivation make CRISPR-Cas systems ideal for neutralizing antibiotic-resistant and rapidly evolving pathogens. In principle, such CRISPR-Cas systems can also be employed as antifungals. However, the conventional CRISPR-based antimicrobial design that mostly relies on the high lethality of Cas-induced DNA double-strand-breaks (DSBs) faces an additional barrier to neutralizing fungal pathogens because robust eukaryotic DNA repair mechanisms can easily counteract DSB activity, rendering Cas enzymes only mildly toxic, if at all (15). Due to this limitation, a new strategy of co-targeting essential genes along with robust repair defensive genes and ensuring their complete or near-complete knockout is vital to achieve efficacious CRISPR-Cas antifungals but is currently unexplored. Since the cataloged diversity of preferred repair mechanisms shows a dependence on growth stages, systematic characterization of essential and repair interactions is critical for the effective design of CRISPR-Cas antifungals but is challenging due to a large combinatorial search of optimal co-targets, especially when facing non-model fungal pathogens. Therefore, the use of a model organism to first understand the significance of this relationship is essential for formulating effective CRISPR-Cas antifungals.

The yeast *Saccharomyces cerevisiae* is a widely studied model organism for understanding fungal genetics and biology (16). Due to its highly curated genome, the essentiality of its genes is well documented and much is known of its robust DNA repair pathways and growth cycles, which exhibit many similarities to the clinically relevant pathogenic yeasts of the *Candida* genus (17–19). Even though *S. cerevisiae* is generally considered to be safe, some strains are known to be opportunistic human pathogens that can be deadly to immunocompromised individuals (20). Thus, the established genetics and clinical relevance make *S. cerevisiae* an ideal model for studying the effectiveness of essential and defensive gene co-targeting as a CRISPR-Cas antifungal strategy.

In this study, we investigated promising CRISPR-Cas sequences for use as antifungals by screening multiple target sites across essential genes that are important for cell survival in *S. cerevisiae*. We demonstrated the enhanced overall lethality of CRISPR-Cas antifungals by co-targeting essential genes with DNA repair defensive genes that removed the ability of the organism to effectively repair the DSB sites. Through modulations that affect cell growth kinetics such as solid versus liquid growth media and cell inoculation, we identified the control of the abundant expression of CRISPR-Cas systems is critical to enhance potency of CRISPR-Cas antifungals while minimizing antifungal resistance. Overall, this study presents a novel design principle for formulating effective CRISPR-Cas antifungals and provides mechanistic insights into the potential rise of antifungal resistance.

## Results and Discussion

### Co-targeting essential and defensive gene machinery is more effective for CRISPR-Cas antifungal formulation

The robust DNA repair mechanisms of eukaryotes present a significant hindrance to the development of CRISPR-Cas antifungals (Fig. 1A). We hypothesized that co-targeting both essential and defensive gene machinery increases the potency of CRISPR-Cas antifungals (Figs. 1B). To test this hypothesis, we investigated the capability of multiplexing gRNAs targeting both essential and defensive gene machinery in *S. cerevisiae*. We began by choosing a set of 3 DNA repair defensive genes—RAD51, RAD52, and LIF1—that are key elements to either homology-directed repair (HDR) or non-homologous end joining (NHEJ) pathways and a set of 23 essential genes involving replication, transcription, translation and metabolism regulation (SI Appendix, Table S1). We then constructed 55 strains with duet plasmid systems carrying gRNAs targeting 55 essential-defensive (E-D) genes on one plasmid and Cas9 on a second plasmid (SI Appendix, Tables S2, S3). As a control, we also constructed 30 strains carrying gRNAs targeting 23 essential (E), 3 defensive (D), and 3 defensive-defensive (D-D) genes. CRISPR-Cas antifungals in these strains were programmed for activation by galactose induction of Cas9 expression. We next characterized the potency of the CRISPR-Cas antifungals by both a serial dilution growth assay on solid media and in liquid culture where cells would not grow in the presence of galactose if the CRISPR-Cas antifungal treatment were effective.

**Figure 1:**
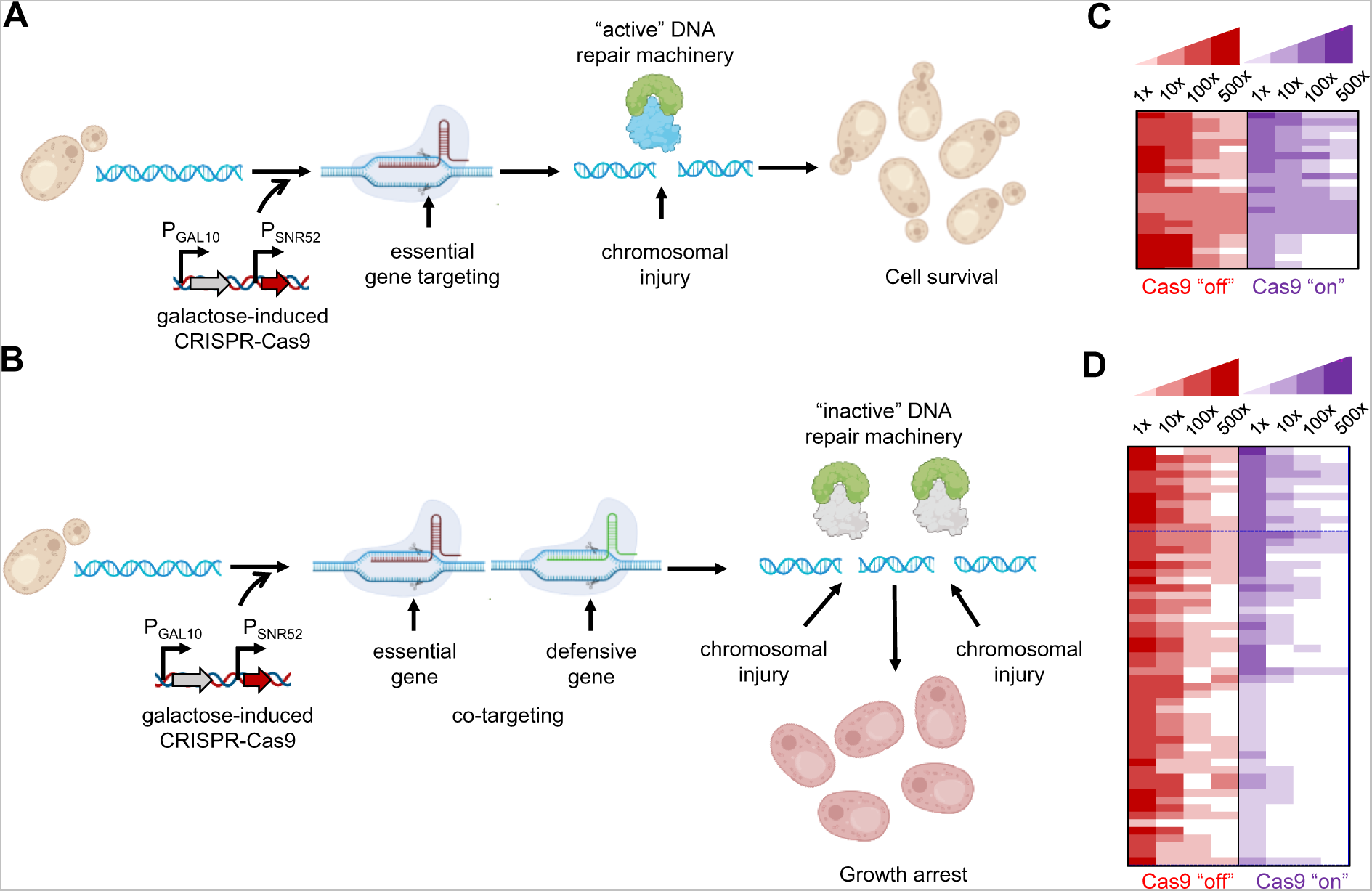
Design of CRISPR-Cas antifungals. **(A, B)** CRISPR-Cas antifungal formulation (**A**) with essential gene targeting alone and **(B)** with essential and defensive gene co-targeting. Targeting a single essential gene locus alone is not effective to eliminate cell survival due to active DNA repair machinery. Co-targeting parts of the DNA repair response reduce the cell’s capability to repair legions at essential and defensive sites, increasing lethality via essential gene knockout and DSB-induced apoptosis. **(C, D)** Heat maps show the effectiveness of **(C)** essential gene targeting and **(D)** essential and defensive gene co-targeting. Each heat map represents serial dilution (1x, 10x, 100x, and 500x) of cells that were treated with different CRISPR-Cas antifungals, spotted on galactose plates (Cas9 “on”, test) and glucose plates (Cas9 “off”, control), and incubated for 48 hours at 28°C. Non-induced strains grown on glucose plates are indicated in red while induced Cas9 strains grown on galactose pads are indicated in purple. Larger colony size is represented by darker coloration. Note that panels **B** and **D** show a representative list of characterized strains; the complete list can be found in SI Appendix, Fig. S1.

In the case of gRNAs targeting single essential or defensive gene loci, we found most strains still exhibited a viable phenotype (Fig. 1C; SI Appendix, Fig. S1) likely because the robust DNA repair machinery of *S. cerevisiae* repaired any breaks at these loci, resulting in only mild toxicity. In contrast, co-targeting of essential and defensive genes proved more effective at inhibiting growth (Fig. 1D; SI Appendix, Fig. S1). As a control, we also found that dual or multiplex targeting essential genes further increased toxicity but fell short of the efficacy found by co-targeting essential and defensive gene machinery simultaneously, which leaves the cell unable to reliably repair a break at the essential gene sites (SI Appendix, Fig. S1). We identified 17 strains (Fig. 2), including Vip47 (harboring LIF1, DNA19), Vip48 (LIF, ACC2), Vip52 (LIF1, RPL10E), Vip55 (LIF1, RPS13B), Vip56 (RAD51, DNA43), Vip57 (RAD51, ISP45), Vip59 (RAD51, RPL7), Vip60 (RAD51, RPS21), Vip69 (RAD53, RPL17), Vip73 (RAD52, RPL7), Vip74 (LIF1, RPL1), Vip76 (RAD52, LIG1), Vip81 (RAD51, RPS4), Vip90 (RAD55, RPS4), Vip93 (LIF1, RSP42), and Vip94 (RAD52, RPO22), that exhibited more effective potency of CRISPR-Cas antifungals using the serial dilution growth assay on solid media. Overall, co-targeting both essential and defensive gene targets is a promising design principle for formulating the most effective CRISPR-Cas antifungals.

**Figure 2.**
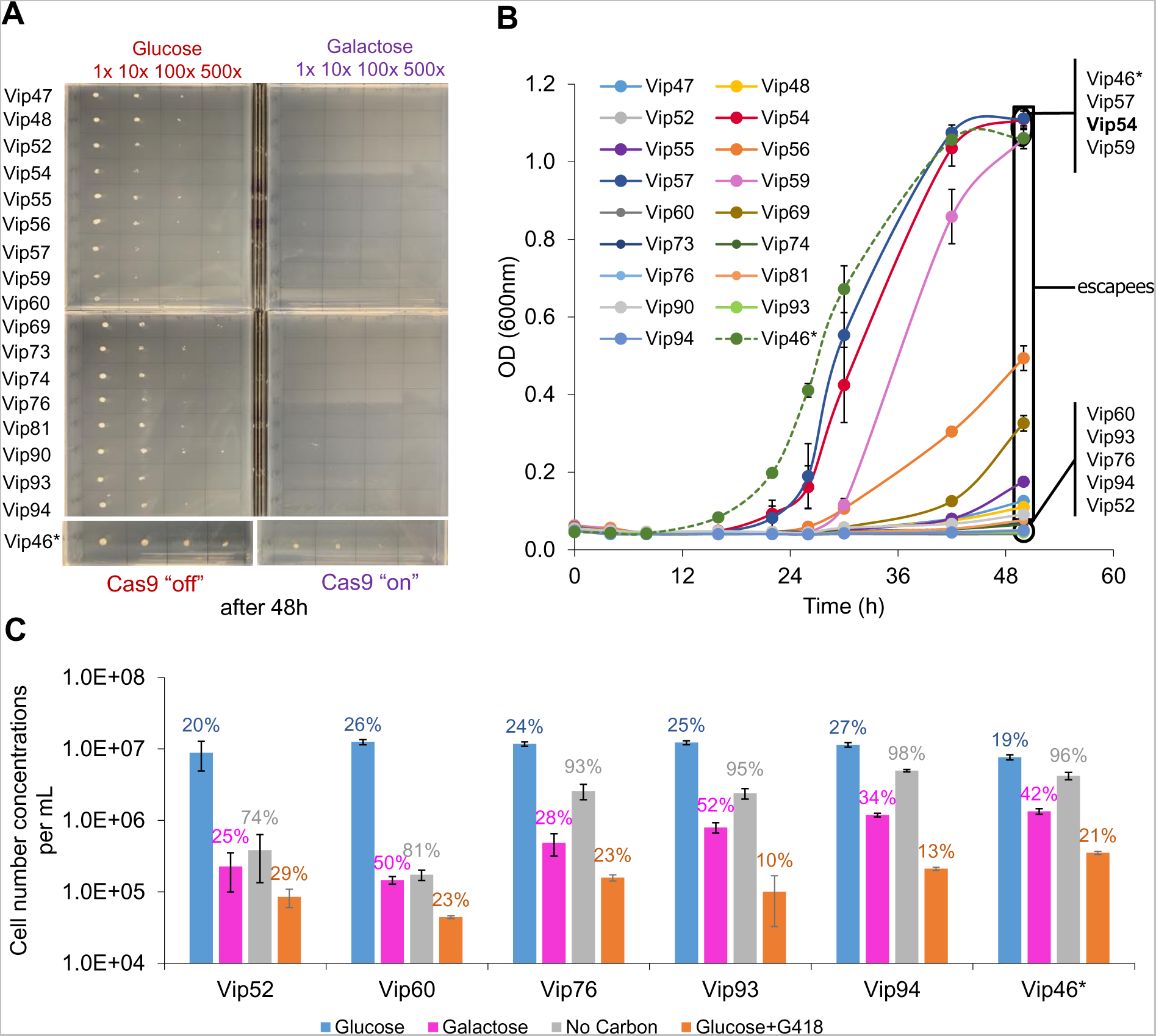
Potency of CRISPR-Cas antifungals and potential rise of antifungal resistance. **(A)** Slower cell growth on solid media did not induce antifungal resistance. All selected strains carrying the top performing CRISPR-Cas antifungals exhibited loss of growth on galactose (Cas9 “on”) plate. **(B)** Faster cell growth on liquid media escaped CRISPR-Cas antifungal treatment. Three strains (Vip54, Vip57, and Vip59) exhibited a normal growth phenotype as compared to the control strain (Vip46*); nine strains (Vip47, Vip48, Vip55, Vip56, Vip69, Vip73, Vip74, Vip81, and Vip90) showed a growth lag but started to enter exponential growth after 48-hour outgrowth; and the remaining five strains (Vip52, Vip60, Vip76, Vip93, and Vip94) exhibited growth arrest. **(C)** Comparison of potency between the top CRISPR-Cas antifungals and antibiotic Geneticin (G418). Cell growth on glucose served as a negative control with inactive CRISPR-Cas systems while cell characterization without a carbon source served as a positive control. In panels **B-C**, each value is a mean ± 1 standard deviation (n ≥ 2).

### Fast cell growth kinetics induced CRISPR-Cas antifungal resistance

We next asked whether cell growth kinetics affected the potency of CRISPR-Cas antifungals. We hypothesized that the fast-growing cells could escape the treatment, propagate, and eventually dominate the entire culture. To test the hypothesis, we characterized growth of the 17 promising strains carrying the CRISPR-Cas antifungals on solid and liquid media, where cells grow faster in liquid media than on solid media. Consistent with the initial screening, none of the strains grew on solid media when CRISPR-Cas antifungals were activated (Fig. 2A). However, only five out of the 17 strains, including Vip52, Vip60, Vip76, Vip93, and Vip94, exhibited flat growth curves on liquid media (Fig. 2B), suggesting greater efficacy of CRISPR-Cas growth inhibition. The remaining strains were escapees from the CRISPR-Cas antifungal treatment, due to faster growth on liquid than solid media. Using one of the top-performing strains, Vip60 (RAD51, RPS21), in a case study, we found that placement of the induced strain in non-inducing media (glucose as carbon source) after 48 hours was able to recover growth (SI Appendix, Fig. S2). In addition, CRISPR-Cas antifungal in Vip60 was active and potent for at least two rounds of culture transfer in liquid media.

To evaluate the potency of CRISPR-Cas antifungals in detail, we compared the cell viability of five high-performing strains under four different growth conditions, including no treatment (normal growth on glucose, negative control), carbon starvation (positive control), geneticin treatment (G418 antibiotic, test case), and CRISPR-Cas antifungals (test case), (Fig. 2C). Under the normal growth condition on glucose without any treatment, all strains grew quickly but exited exponential growth phase after 48 hours with a high count of viable cells but a low percentage of cell viability (19-27%). When subjected to carbon starvation, all strains experienced growth inhibition with a high percentage of cell viability (74-98%) after 48 hours, implying the cells were mostly dormant. In the presence of antibiotic treatment, cells exhibited not only growth inhibition but also a low percentage of cell viability (10-29%). As compared to the carbon starvation scenario, all strains under CRISPR-Cas antifungal treatment showed more severe growth inhibition with a lower percentage of cell viability (25-52%). CRISPR-Cas antifungals in both Vip52 and Vip60 strains exhibited high potency like G418 antibiotic treatment with low cell count and low percentage of cell viability while others mostly showed dormant phenotypes (low cell count and high percentage of cell viability) like under carbon starvation. In our design, Vip52 co-targets the defensive gene LIF1 (ligase interacting factor mediating NHEJ in DNA double-strand break repair) and the essential gene RPL10E (ribosomal protein of 60S unit), while Vip60 co-targets the defensive gene RAD51 (radiation sensitive protein mediating HDR in DNA double-strand break) and the essential gene RPS21 (ribosomal protein of 40S unit). Even though Vip52 and Vip60 are designed to target different gene sets, their potency as antifungals signifies the importance of the essential and defensive gene co-targeting strategy.

Taken altogether, CRISPR-Cas systems can be potentially used as novel antifungals by co-targeting defensive and essential genes. Identifying optimal combinations of the essential and defensive gene targets that outpace the fast cell growth kinetics and cause cell death rather than cellular dormancy will be critical in formulating the effective CRISPR-Cas antifungals.

### High cell density caused a more pronounced community escape of CRISPR-Cas antifungal treatment

To further understand the robustness of growth suppression in the top-performing strains, we investigated the effect of high cell density inoculation on the potency of CRISPR-Cas antifungals. We differentially seeded cultures with either a low inoculation (0.05 OD, 5.5×10^5^ cells/mL, as conducted in previous experiments) or a high inoculation cell density (0.2 OD, 2.5×10^6^ cells/mL). Without carbon starvation, the increase in cell inoculation resulted in a pronounced increase in growth rates with a shorter lag phase in all media types (Figs. 3A-3F). While the escape phenomenon (or CRISPR-Cas antifungal resistance) dominated in high cell density inoculation, it was more significant for the antibiotic treatment than the CRISPR-Cas antifungal treatment. Consistent with growth characterization in solid versus liquid media, the CRISPR-Cas antifungal activities of Vip52 and Vip60 also exhibited the best performance with high cell inoculation scenario among the characterized strains (Fig. 3G). The resistance to G418 in high cell inoculation, but not in low cell inoculation, was likely because the greater number of cells produced a greater effective concentration of aminoglycoside 3’-phosphotransferases and hence provided the innate resistance via degradation of the antibiotic (1). This broad escape phenomenon induced by higher initial cell concentration—and therefore higher loading of enzymes responsible for counteracting antimicrobial activity—suggests that ongoing kinetics of the culture is also a major driving factor of the escape from CRISPR-Cas antifungal treatment, in contrast to an initial inhibition/inefficiency of the machinery that eventually allows for escaped cells to proliferate. This escape phenomenon was more pronounced when CRISPR-Cas antifungals were less effective as seen in the non-top-performing strains (SI Appendix, Fig. S3). Overall, faster growth kinetics in high cell inoculation reduced the potency of CRISPR-Cas antifungals and caused pronounced community escape as observed in solid versus liquid media. The differential inoculation results underscore the similarity in escape profiles between traditional small-molecule antibiotics and CRISPR-Cas antifungals, requiring deeper investigation into the roles of cellular repair machinery, stress response mechanisms, and effective CRISPR-Cas systems (Cas enzymes and gRNAs) to this phenotype.

**Figure 3.**
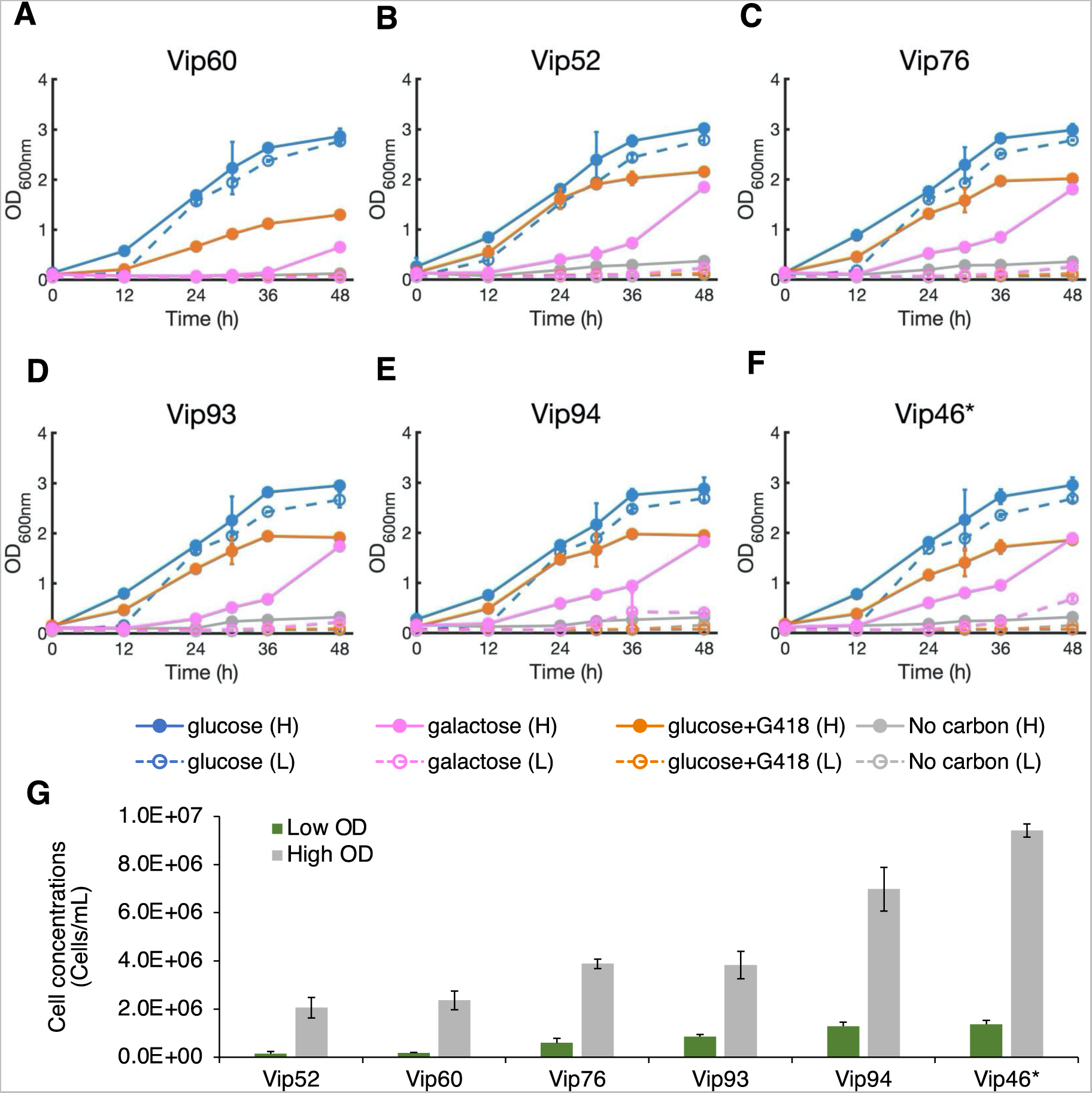
Higher cell inoculation is more prone to CRISPR-Cas antifungal resistance. **(A-F)** Effect of cell inoculation concentrations on the potency of CRISPR-Cas antifungals. Five strains (Vip52, Vip60, Vip76, Vip93, and Vip94) harboring the most effective antifungal designs and one control strain (Vip46*) were investigated with an initial OD of either 0.05 or 0.2. Higher concentration of cells at the outset of the experiment overwhelmed the population by decreasing lag time, thereby increasing both antibiotic and CRISPR-Cas antifungal resistance. **(G)** Viable cell count at 36-hour outgrowth confirmed the increase in viable cells in the high inoculation scenario. In panels **A-G**, each value is a mean ± 1 standard deviation (n ≥ 2).

### Genotypes of escape population contained small mutation frequency in targeted genes and CRISPR-Cas sequences

To better understand how cells escaped the CRISPR-Cas antifungal treatment, we performed both amplicon and plasmid sequencing for representative strains harboring single and co-targeting CRISPR-Cas antifungals using a combination of Illumina (MiSeq) and Oxford Nanopore sequencing (Fig. 4A). Our results showed that escape populations were primarily comprised of cells that have avoided mutation to the targeted genes (Figs. 4B, 4C) as well as any mutations on the plasmid sequence harboring CRISPR-Cas9 sequences (Figs. 4D, 4E). Illumina sequencing of approximately 1 kb PCR-amplified regions around the Cas9 target site on the genome revealed mutation frequencies of less than 5% for essential genes and 8%, 6%, and 7% for RAD51, RAD52, and LIF1, respectively (Fig. 4B). These mutations were dominated by large deletions (> 20 bp) destroying the target sites and hence the coding sequences of the target genes, implying gene knockouts (Figs. 4C; SI Appendix, Fig. S4). However, as these mutated sequences represent a small minority of the total population, it suggests that the dominance of HDR over NHEJ repairs prevents significant edits even at nonessential gene loci. The rapid division of cells with shortened lag phases provides a considerable number of templates for HDR machinery to overcome the CRISPR-Cas rate of double-strand breaks, as observed for cultures growing in solid versus liquid media (Fig. 2) and high cell inoculation (Fig. 3).

**Figure 4.**
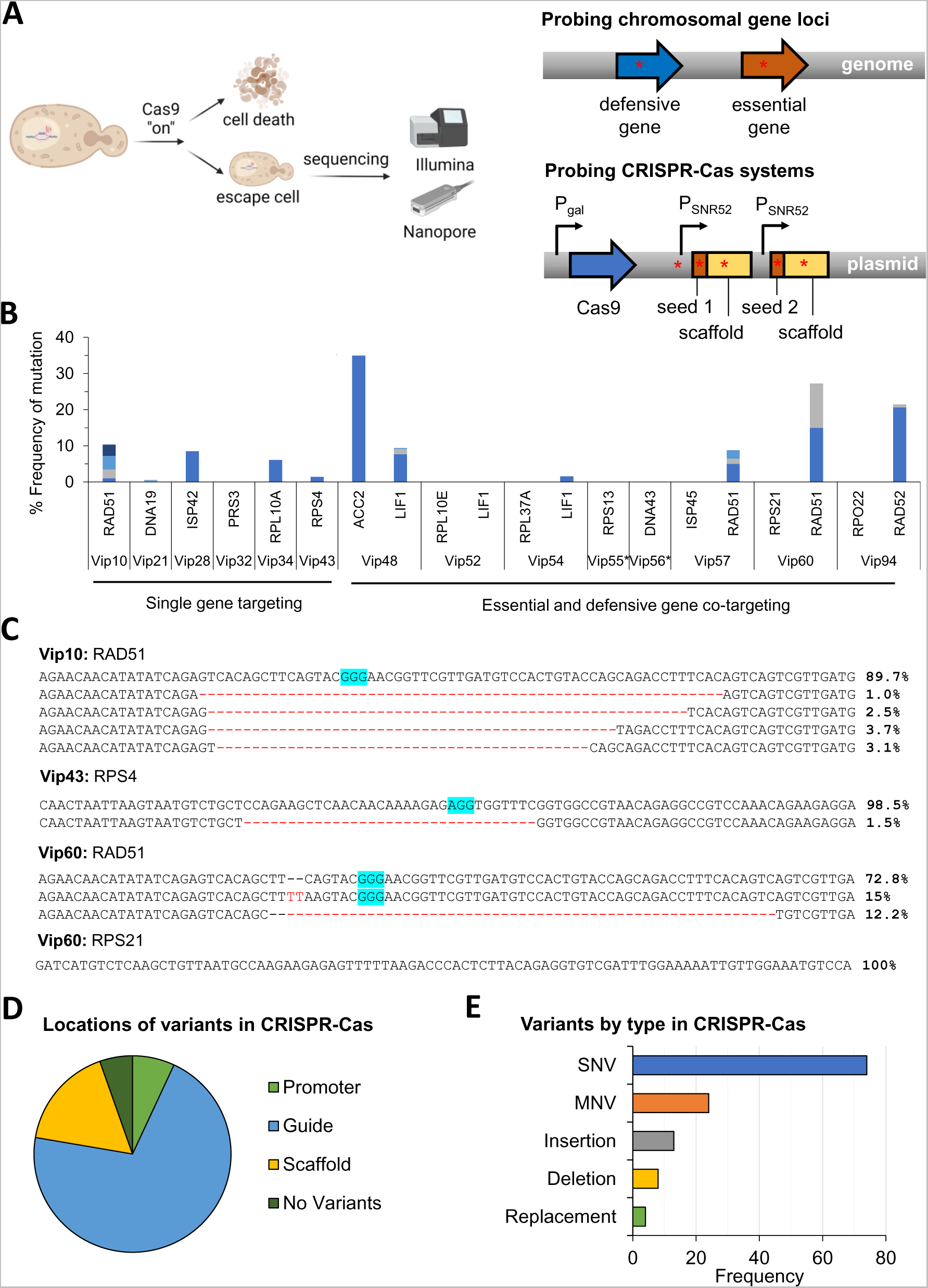
Identification of mutations from representative single- and co-targeted strains that escape from the CRISPR-Cas antifungal treatment. **(A)** Experimental workflow for sequencing. **(B)** Genetic mutation frequency of the targeted chromosomal genes. Different colored stacked bars represent different mutations presented in Fig. 4C and SI Appendix, Fig. S4. **(C)** Mutated sequences for the representative strains Vip10, Vip43, and Vip60. PAM sequences are highlighted in blue. Single targeted strains show low levels of mutation with the exception of the defensive genes, in particular RAD51. As seen in the sequence alignments provided for Vip10, Vip43, and Vip60, mutants are dominated by large (>20 bp) deletions, although mutation frequency is low and heterogeneous. Mutated sequences for other characterized strains were presented in SI Appendix, Fig. S4. **(D)** Locations of mutations in the gRNA cassettes on extracted gRNA plasmids. **(E)** Variants by type in gRNA cassettes.

In the co-targeted strains, targeted sequencing revealed that the defensive gene loci (i.e., LIF1, RAD51, and RAD52) still experienced a small amount of mutation while mutations in essential genes were still undetectable, except for a small deletion (2 bp) in the ACC2 gene in the Vip48 strain (Figs. 4B; SI Appendix, Fig. S4). The results in the co-targeted strains are consistent with the single-targeted strains, implying that the surviving populations, while having an active CRISPR-Cas system, were able to repair their breaks in essential genes with the accurate HDR repair method. Additionally, because the defensive genes did not experience a full knockout, a lowered but sufficient amount of repair proteins were expressed for DNA damage repair of essential loci to escape the CRISPR-Cas antifungal treatment.

Oxford Nanopore resequencing of the Cas9-bearing plasmids extracted from the 17 high-performing strains (average sequencing depth of 20,000x) revealed no detectable rearrangements on the plasmid and no nucleotide variants over a 1% threshold, indicating that, unlike in bacterial pathogens, Cas9 mutation is not the dominant pathway to escape. Illumina amplicon sequencing of the guide cassette from gRNA plasmids revealed high frequencies of mutation at this site of the plasmids (Figs. 4D, 4E). Mutations occurred at various sites along the gRNA operon including the promoter (7%), scaffold (17%), and target sequence (71%) (Fig. 4D). Single nucleotide variants (SNVs) were the most common (60%) but the remaining 40% of mutations were made up of indels and other multi-nucleotide variants (MNVs), which lead to greater loss-of-function (Fig. 4E). This sequencing result—consistent with previous studies in other species (21–24)—indicates that the primary route of CRISPR-Cas antimicrobial evasion is the mutation of the gRNA operon on the delivered plasmid. This study shows that *Saccharomyces* and eukaryotes more broadly also exhibit increased resistance over bacteria due to the robust and redundant DNA repair mechanisms.

Taken together, the competition among kinetics of cell growth, DNA repair, and CRISPR-Cas activity is a key factor controlling antifungal effectiveness. Increasing the efficiency of gene knockouts through the selection of superior Cas proteins and gRNAs in conjunction with more effective ways to reduce the DNA repair pathway expression will be critical to deploy CRISPR-Cas systems as an antifungal therapy.

## Conclusions

The development of novel antifungals is a critical area of concern for public health, agriculture, and the environment. This study presented the design principles and identified key factors and areas of improvement for effective CRISPR-Cas antifungal formulation. Importantly, the robust DNA repair mechanisms encountered in eukaryotic pathogens must be addressed by means of direct targeting or utilization as a “Trojan Horse” to counteract their role in directing cells to a persister phenotype. This study underscores the importance of such a co-targeting approach of essential and defensive genes to increase CRISPR-Cas antifungal activity. In addition, informed selection of these defensive genes is needed based on the pathogen in question, as certain pathways are favored in different species. Future work should focus on evaluating the temporal dimension to determine means of preventing escaped persister cells from re-colonizing a host. Furthermore, the delivery of CRISPR-Cas systems to fungal pathogens *in vivo* must be addressed for these systems to function as a viable antifungal strategy.

## Materials and Methods

### Strains and plasmids

A complete list of plasmids, strains, and primers of this study are presented in Tables S2, S3, and S4 of SI Appendix, respectively. The plasmid vectors p415-GalL-Cas9- CYC1t (Addgene plasmid # 43804) and p426-SNR52p-gRNA.CAN1.Y-SUP4t (Addgene plasmid # 43803) were a gift from George Church and were used as the backbone plasmids for all studies (25). All gRNAs were designed using CASPER (26, 27).The p426 derived vectors containing the gRNA sequences were built via Phusion PCR in a 40 µL reaction using the p426 plasmid as a template, gRNA Gibson primers with homology for the template and the insert, and a generic primer with homology for the plasmid (AT_gRNA_BB_F or AT_gRNA_BB_R). Inserts were constructed using the same template and complementary primers to the backbone (i.e., one unique to a gRNA and one generic). Linker sequences were used to construct an insert with one gRNA sequence followed by the SUP4t terminator, a short spacing sequence, and then another SNR52 promoter with the second gRNA sequence on the end. These inserts were inserted into the p426 backbone using the same Gibson assembly protocol described (28).

Transformations were performed with Top10 *E. coli* using 5 µL of reacted Gibson Assembly. Upon addition, cells were left on ice for 3 hours followed by a 45-seconds heat shock at 42°C. After heat shock, cells were inoculated in 1 mL of lysogeny broth (LB) and shaken for 1 hour at 37°C. Cultures were spun down and concentrated for plating on 100 mg/L Amp plates, where they grew overnight at 37°C. Transformants were confirmed by colony PCR and plasmids were confirmed by Sanger sequencing following plasmid extraction. Approximately 1 μg of each confirmed plasmid with the Cas9 carrying plasmid were transformed into *S. cerevisiae* BY4741 using electroporation (2.1 kV, 200 Ω, 25 µF).

### Media and culturing conditions

The parent strain used in this study was *S. cerevisiae* BY4741. Strains carrying the Cas9 and gRNA plasmids were cultured in selective media with the appropriate double auxotrophy (SC-Ura^-^Leu^-^), the desired carbon source (20 g/L glucose or 20 g/L galactose for induction of Cas9). For agar plate serial dilution assay, strains were cultured in SC-Ura^-^Leu^-^ plus glucose media overnight at 28°C until exponential phase was reached. They were then washed once with PBS, diluted 10x, 100x, and 500x in PBS, spotted 1 μL of sample on the agar plate, and incubated at 28°C for 48-72 hours. For growth assays in broth, culture preparation followed the same incubation followed by PBS wash. Cultures were then diluted with culture media (SC-Ura^-^Leu^-^ plus carbon source) to the appropriate starting OD (0.05 or 0.2). Strains were cultured in a Duetz plate with a working volume of 500 μL in a maxQ 6000 shaker (Thermo Fisher) set at 28°C and 400 rpm. OD measurements were taken with a Bio-Tek plate reader.

### Flow cytometry

To perform viability assessment with flow cytometry, samples were taken and diluted to within a range of OD 0.1-1.0 to ensure accurate counting by the flow cytometer. Samples were then further diluted 20x to a total volume of 200 µL by the Guava ViaCount staining solution (EMD Millipore #4000-0041), mixed by pipetting, and allowed to sit in the dark for 5 minutes. Samples were taken in a 96-well round bottom plate and placed in the Guava EasyCyte 6HT flow cytometer. Gating voltages were calibrated on a 1:1 mixture of an exponential phase BY4741 culture and a 20-second microwaved sample of the same strain.

### Sequencing to identify mutated sequences

A deep-hybrid sequencing using Illumina MiSeq and Oxford Nanopore was performed to examine both the target chromosomal loci and the plasmids carrying CRISPR-Cas from escaped strains after the CRISPR-Cas antifungal treatment. Briefly, plasmids and genomic DNA from selected strains were extracted 48 hours after galactose induction. For examining the target chromosomal loci, about 1 kb around each target locus was PCR amplified from 100 ng of extracted genomic DNA with Phusion Hot Start II DNA Polymerase (Thermo Fisher) using the locus-specific primers found in SI Appendix, Table S4. Amplicons were purified using the Omega Biotek E.Z.N.A Cycle Pure Kit (SKU: D6492-01) and pooled in equimolar ratios. For investigating gRNA regions from extracted plasmids, the promoter plus guide RNA region was PCR amplified using primers AT-gRNA-swap_F and AT-gRNA-swap_R, purified, and pooled in equimolar ratios.

For Illumina sequencing of the target chromosomal loci and plasmid gRNA regions, amplicons were further prepared as genomes using the Nextera XT library preparation kit and evaluated on a bioanalyzer for quality control. Pools were then all combined and diluted to 4 nM. Final products were diluted to a final loading concentration of 4 pM, pooled with 20% of 10 pM PhiX, and loaded on a Version 3 flow cell reading 275 bp, paired-end, on the Illumina MiSeq at the University of Tennessee Genomics Core.

For Oxford Nanopore sequencing of target chromosomal loci and the extracted Cas9 and gRNA plasmids, amplicons and plasmids were prepared using the LSK109 ligation sequencing kit with the native barcoding kit (EXP-NBD104) and sequenced on a MinION R9.4.1 flow cell with an average sequencing depth of 20,000x. Sequencing data was imported and analyzed in the Qiagen CLC Genomics Workbench 20.0.4. Indels and rearrangements in target loci were analyzed using the InDels and Structural Variants tool. Variants in the Cas9 plasmids and gRNA regions were analyzed using the Low Frequency Variant Detection tool with a minimum frequency threshold of 1%.

## Data, Materials, and Software Availability

All data of this study are included in the article and SI Appendix.

## Acknowledgements

The research was financially supported by the DARPA YFA award and Director Fellowship (D17AP00023). The authors would like to thank Dr. Renee Wegrzyn and her DARPA team for useful discussion during the course of the project. The views, opinions, and/or findings contained in this article are those of the authors and should not be interpreted as representing the official views or policies, either expressed or implied, of the funding agency.

## Author contributions

CTT conceived the research study. CTT and BJM designed the research. BJM, XZ, and JCC performed research; BJM, CC, and CTT analyzed data; and BJM, CC, and CTT wrote the paper.

## Supplementary Information

**Figure S1:** Screening potency of CRISPR-Cas antifungals. Vip strains that harbor gRNAs targeting 23 essential (E), 3 defensive (D), 4 defensive-defensive (D-D), and 56 essential-defensive (E-D) genes were characterized for growth using the serial dilution (1x, 10x, 100x, 500x) growth assay on solid media after 48 hours. CRISPR-Cas9 systems were active in the presence of galactose but were inactive in the presence of glucose (negative control). Growth phenotypes were ranked from 0 to 4, with 0 indicating no growth and 4 indicating a healthy colony and used to generate the heat map. A subset of the top 17 strains whose names are shown in red exhibited high potency of CRISPR-Cas antifungals, co-targeting both essential and defensive genes. These strains were selected based on the criteria that they grew well on glucose plates but could not grow on galactose plates after 48 h. Biological replicates: n = 2.

**Figure S2:** Effect of growth adaptation in liquid cultures on potency of CRISPR-Cas antifungal (Vip60). **(A)** Seed preparation**. (B)** Round 1 of CRISPR-Cas antifungal treatment. **(C)** Round 2 of CRISPR-Cas treatment. Experiments were performed in biological triplicates.

**Figure S3.** Effect of cell inoculation density on escapee proliferation in non-top performing Vip strains. Strains were investigated with an inoculation OD of either 0.05 (L) or 0.2 (H). The escape phenomenon was more pronounced for less effective Vip strains at both high and low inoculation densities.

**Figure S4:** Sequence alignments from targeted amplicon sequencing of representative single- and co-targeted strains. PAM sequences are highlighted in blue (sense) and pink (antisense).

**Table S1:** Gene annotation. Sources: Saccharomyces Genome database. See https://www.yeastgenome.org/

**Table S2:** List of plasmids. Abbreviations: E, essential; D, defensive. See Table S1 for gene annotation.

**Table S3:** List of strains. See Table S1 for gene annotation.

**Table S4:** List of primers.

